# Detection and Polarization of Introgression in a Five-taxon Phylogeny

**DOI:** 10.1101/004689

**Authors:** James B. Pease, Matthew W. Hahn

## Abstract

In clades of closely related taxa, discordant genealogies due to incomplete lineage sorting (ILS) can complicate the detection of introgression. The *D*-statistic (a.k.a. the ABBA/BABA test) was proposed to infer introgression in the presence of ILS for a four-taxon clade. However, the original *D*-statistic cannot be directly applied to a symmetric five-taxon phylogeny, and the direction of introgression cannot be inferred for any tree topology. Here we explore the issues associated with previous methods for adapting the *D*-statistic to a larger tree topology, and propose new “*D*_*FOIL*_” tests to infer both the taxa involved in and the direction of introgressions for a symmetric five-taxon phylogeny. Using theory and simulations, we find that previous modifications of the *D*-statistic to five-taxon phylogenies incorrectly identify both the pairs of taxa exchanging migrants as well as the direction of introgression. The *D*_*FOIL*_ statistics are shown to overcome this deficiency and to correctly determine the direction of introgressions. The *D*_*FOIL*_ tests are relatively simple and computationally inexpensive to calculate, and can be easily applied to various phylogenomic datasets. In addition, our general approach to the problem of introgression detection could be adapted to larger tree topologies and other models of sequence evolution.

---

In phylogenomic analyses of closely related taxa, conflicting phylogenetic signals among loci are a common occurrence. Discordant genealogies (ones that disagree with each other and possibly the true species topology) represent both a challenge in determining the true species phylogeny and a potential source of additional information about a clade’s evolutionary history (Maddison 1997; Degnan and Rosenberg 2009; Edwards 2009). Rapid successive speciation events at any time before the present can lead to discordant genealogies via incomplete lineage sorting (ILS), where two lineages fail to coalesce within a population, making it possible for either lineage to coalesce first with a less related population (Hudson 1983; Tajima 1983; Pamilo and Nei 1988). Discordant genealogies of closely related species can also occur through various forms of hybridization, ranging in scope from the introgression of alleles between existing species to the formation of new hybrid species (Currat et al. 2008; Twyford and Ennos 2012). Even though ILS and introgression/hybridization cause discordant gene trees, the difference in their effects on tree topologies allows them to be disentangled. While several methods for the detection of hybrid speciation in clades with ILS have been previously proposed (Sang and Zhong 2000; Meng and Kubatko 2009; Yu et al. 2012; Yu et al. 2013), here we focus on the detection of introgression between distinct, non-hybrid species in a clade where ILS is present.

Several methods for distinguishing ILS from introgression in a four-taxon phylogeny (three species and an out-group; Fig. 1a) have been proposed previously. The *D*-statistic (a.k.a. the “ABBA/BABA test”) looks for an imbalance in the relative frequency of two minor, discordant gene trees (Green et al. 2010; Durand et al. 2011). Given a consensus phylogeny, ILS should produce the two minor topologies with equal frequency. However, introgression causes an imbalance toward a closer relationship between the two taxa exchanging alleles. Therefore, a statistically significant imbalance toward one discordant topology (indicated by the allele pattern ABBA or BABA) indicates that introgression has occurred.

The *4sp* algorithm (Garrigan et al. 2012) also uses the relative frequencies of allelic state patterns to determine regions of introgression. This algorithm uses a maximum-likelihood approach to estimate global parameters of the species tree, then calculates a local likelihood of introgression for each region of the genome (Garrigan et al. 2012). Joly et al. (2009; see also Joly 2012) use a similar method to look for loci that have sequences with unusually low genetic distances. A significantly lower minimum sequence distance for a pair of taxa indicates introgression at specific loci. Both of these methods, and the *D*-statistic, allow for fine-scale analysis of localized introgression when genomic spatial information is available (i.e., large chromosomal alignments).

When expanding phylogenetic tests for ILS and introgression beyond the simple four-taxon case, there are several challenges. In the presence of ILS, a four-taxon, rooted phylogeny (Fig. 1a) has only three different possible gene tree topologies. For a five-taxon species phylogeny (Fig. 1b) in the presence of ILS there are fifteen possible gene tree topologies both symmetric (Fig. 1c) and asymmetric (Fig. 1d). In addition to more possible gene trees, there are an increased number of possible introgression donor-recipient pairs, and the probability distribution of possible gene trees must also be considered (Degnan and Salter 2005; Degnan and Rosenberg 2006; Twyford and Ennos 2012).

Here, we propose a new set of statistical measures (the *D*_*FOIL*_ tests) that not only overcome the inherent challenges of additional phylogenetic information in five-taxon symmetric phylogenies, but also leverage this added data to infer both the taxa involved in and direction of introgression. We also show how these statistics can be used to infer the occurrence of introgression along ancestral branches, and present results that suggest a previous modification of the *D*-statistic (the “Partitioned” *D*-statistic; Eaton and Ree 2013) imprecisely identifies the taxa involved in introgression. The dynamics of all these statistics are shown theoretically and by simulation, followed by a discussion of their applications to data.

## Materials and Methods

### The Four-taxon D-statistic

In order to approach testing introgression in a five-taxon phylogeny, we first describe the four-taxon case. (Note that in much of the literature on the multispecies coalescent these are referred to as three-taxon and four-taxon trees, where the outgroup is not counted. For consistency with previous work on the *D*-statistic, we do not use this terminology here.) The four-taxon *D*-statistic for introgression was formalized to test ancestral admixture between human and Neanderthal populations (Green et al. 2010; Durand et al. 2011). This statistic applies to a four-taxon asymmetric phylogeny with three in-group taxa and an out-group, denoted (((*P*_1_,*P*_2_),*P*_3_),*O*) (Fig. 1a). All sites considered in the alignment of sequences from these taxa must be either mono- or biallelic, with the out-group defining the ancestral state (always named A) relative to the derived state (named B). Allelic state patterns for a given position in the alignment are given in the order *P*_1_*P*_2_*P*_3_*O* (e.g., ABBA; Fig. 1 and 2). Site pattern counts (e.g., *n*_ABBA_) are the raw counts of these site types in a given region of the sequence alignment. The model generally assumes 0 or 1 substitutions at each site over the whole phylogeny, with a negligible number of reverse and convergent substitutions. This model also assumes 0 or 1 introgressions in a region. The true is tree is supported by the patterns BBAA and BBBA, and polyphyletic appearance of B in the discordant site patterns ABBA and BABA is attributed to either ILS or introgression (or both).

The *D*-statistic is calculated as (Green et al. 2010; Durand et al. 2011):

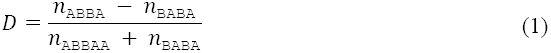

Under ILS and no introgression, the discordant tree patterns ABBA and BABA should occur with equal frequency (Hudson 1983; Tajima 1983; Pamilo and Nei 1988). Introgression between *P*_3_ and either *P*_1_ or *P*_2_ will disproportionately increase the frequency of BABA or ABBA, respectively, since the introgressed pair of taxa should have relatively more shared derived (B) states. Therefore, the four-taxon *D*-statistic is a measure of inequality in the prevalence of site patterns that support the two possible discordant gene tree topologies. Positive values of *D* indicate *P*_2_⇔*P*_3_ and negative values indicate *P*_1_⇔*P*_3_, with values not differing significantly from zero not supporting either introgression (where ⇔ denotes introgression of indeterminate direction and ⇒ denotes a polarized introgression). This approach is not able to detect introgression between the two sister taxa, *P*_1_ and *P*_2_.

### Considerations for Extending the D-statistic Beyond Four Taxa

As the number of taxa in a phylogeny increases, introgression testing becomes increasingly complex due to three factors: (1) the number of introgression donor-recipient pairs increases geometrically, (2) the number of possible gene tree topologies also increases geometrically, and (3) the probability distributions of discordant gene tree topologies under ILS become more complex with both the size and shape of the phylogeny (Rosenberg 2002; Degnan and Salter 2005; Degnan and Rosenberg 2006; Rosenberg 2007). Another practical concern is that the four-taxon *D*-statistic can only provide two answers (positive and or negative), corresponding to the introgressions *P*_1_⇔*P*_3_ and *P*_2_⇔*P*_3_. A single *D*-statistic is sufficient for the four-taxon case, but a larger phylogeny will require a system of multiple tests to distinguish among the (rapidly) increasing number of possible introgressions.

There are multiple perspectives from which the *D*-statistic can be understood. In all interpretations, the *D*-statistic has the null hypothesis that *D*=0 with or without ILS, and with no introgression. From the perspective of site patterns, the *D*-statistic states that the biallelic site patterns ABBA and BABA should be sampled with equal frequency under the null hypothesis (Eq. 1). The site patterns ABBA and BABA are a proxy for the two discordant gene trees (((*P*_2_,*P*_3_),*P*_1_),*O*) and (((*P*_1_,*P*_3_),*P*_2_),*O*), respectively (Fig. 2a, lines ii and iii), each of which is expected to occur with equal frequency. More generally, we can say that the *D*-statistic compares two sets of gene trees—the “left” and “right” terms of the numerator—whose sampling probabilities are expected to be equal given the probability distribution of gene trees for a particular species phylogeny. In a four-taxon phylogeny, the two sets of gene trees are each represented by only a single discordant gene tree (of the two possible discordant topologies). Since the two discordant gene trees are equally likely to be sampled under ILS, there is an equal probability of inferring that *P*_3_ is more closely related to *P*_1_ or *P*_2_, and *D* equals 0 under the null hypothesis of no introgression. Importantly, in the *D*-statistic it is not necessary to calculate the exact probability of sampling either discordant gene tree. Because both discordant trees are due to ILS on the same ancestral branch, both are expected to have equal relative probability regardless of what the absolute probabilities are (Fig. 2a). This feature will be important in designing *D* statistics for five-taxon phylogenies.

Another important aspect of the four-taxon *D*-statistic is that it only uses shared derived states to infer introgression. However, introgression is equally capable of transferring the ancestral state (A) or the derived state (B). For example, consider a four-taxon phylogeny where a substitution A→B has occurred on the internal branch *P*_12_ (Fig. 1a). This would ordinarily lead to the site pattern BBAA. However, if *P*_3_⇒ *P*_1_ introgression occurs, then the A state is transferred resulting in pattern BAAA. This means that when introgression is considered, both ABBA and its “inverse pattern,” BAAA, offer evidence of *P*_3_⇔*P*_1_ introgression, and the same is true for BABA/ABAA and *P*_3_⇔*P*_2_. As long as both terms in the numerator of *D* use these inverse site counts, the null hypothesis of *D*=0 is maintained. However, the inclusion of patterns with a single derived state (i.e. one B) may cause complications in some types of sequence data and in some biological scenarios where substitution rates are unequal among terminal branches of the phylogeny (see Discussion).

From these considerations of the four-taxon *D*-statistic, we can derive four general principles that can be used to test introgression in a five-taxon phylogeny. First, a system of multiple *D*-statistics will be required to distinguish among the larger number of possible donor-recipient combinations of introgression. Second, rather than designing a solution particular to this tree topology as a whole, we can discretize introgression testing into a system of taxon-by-taxon *D*-statistics by examining the relative relationships of a given taxon against two other (appropriately selected) taxa. Third, to maintain the null expectation of *D*=0, these two relative phylogenetic relationships must have an equal sampling probability across the distribution of all possible gene trees. As with the four-taxon statistic, it will not be necessary to calculate the actual probability values as long as gene trees can be selected in equally probable pairs (as illustrated in Figure 2). Finally, inverse patterns (e.g. BABA/ABAA) both indicate the same potential introgressions.

**FIGURE 2.**
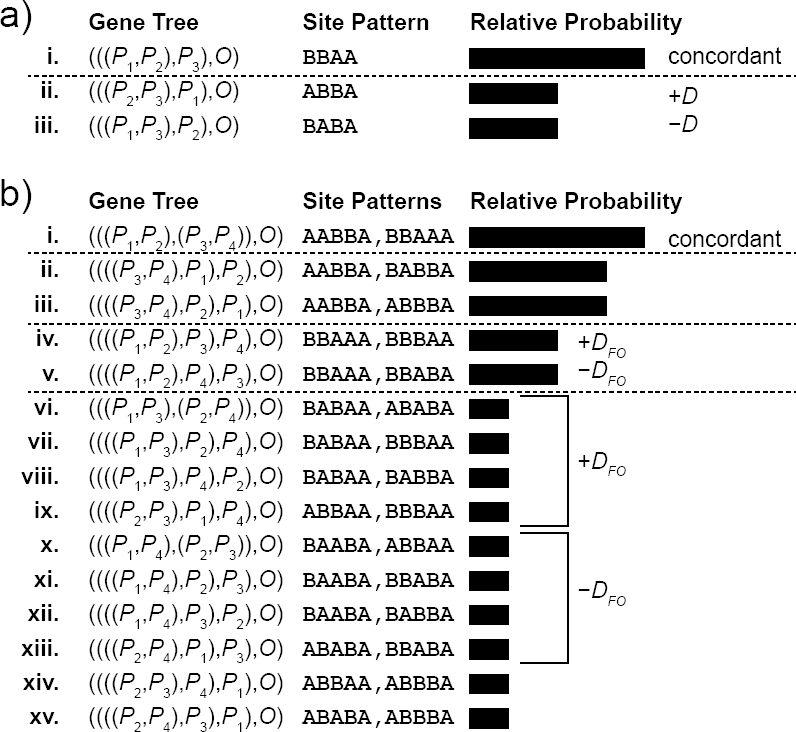
(a) The three possible gene trees for a four-taxon phylogeny, including the gene tree concordant with the species phylogeny (i) and the two discordant gene trees (ii and iii). For each gene tree the biallelic site patterns used in the *D*-statistic are shown. Note that the “left” (+*D*) and “right” (-*D*) terms of the *D*-statistic sample gene trees whose relative probability is equal. (b) The fifteen possible gene trees for a five-taxon phylogeny, including the concordant gene tree (i) and discordant gene trees (ii-xv). The effects of discordant topologies for the *D*_*FO*_ statistic is shown as an example. *D*_*FO*_ samples sets of gene trees for the “left” (+*D*_*FO*_) and “right” (-*D*_*FO*_) terms, whose relative probabilities are equal regardless of their absolute probabilities. The relative values of gene tree pairs where ILS only occurs on a single branch (ii/iii and iv/v) depend on the relative length of the ancestral branches *P*_12_ and *P*_34_.

### The Five-Taxon “Partitioned” D-statistics

We also use these principles to examine the “Partitioned” *D*-statistics previously proposed to infer inter-group introgression (Fig. 3a) in a symmetric five-taxon phylogeny (Eaton and Ree 2013). These statistics are:

**FIGURE 3.**
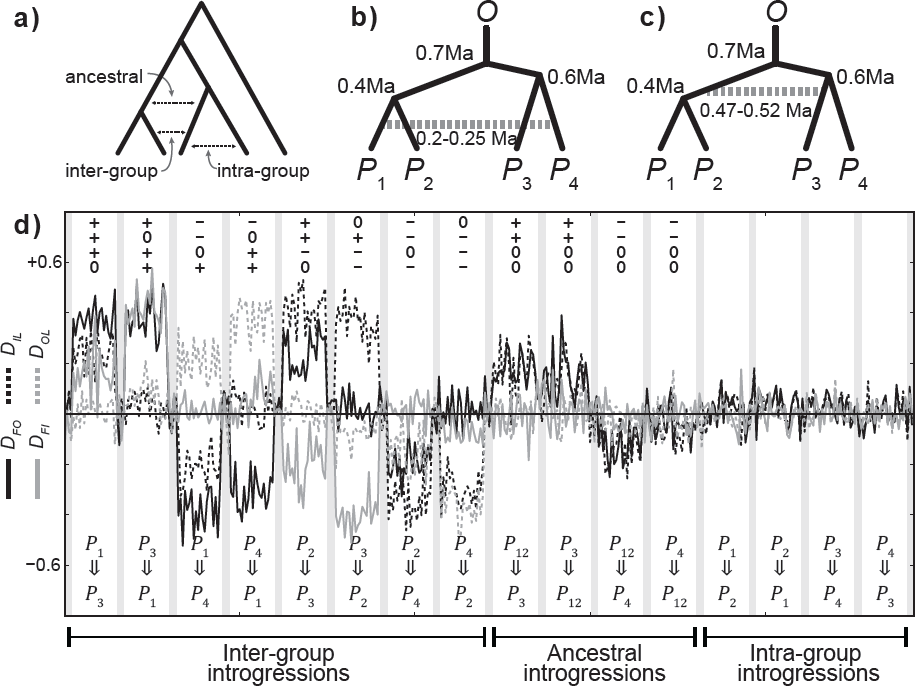
The *D*_*FOIL*_ statistics can detect both the taxa involved in and direction of introgression. (a) In a four-taxon symmetric topology, three classes of introgression can be defined: “intragroup” between taxa in the same subgroup, “inter-group” between taxa in different subgroups, and “ancestral” involving the ancestral population of one subgroup. (b) Topology of tree used for simulations with species divergence times given in terms of millions of years ago (Ma) listed next to their respective nodes and inter-group introgression occurring between 0.20 and 0.25 Ma (dashed line). (c) The same topology showing the time of ancestral introgression between 0.53 and 0.58 Ma. (d) Simulated chromosomes with *n*=500 10 kb regions for each of the sixteen possible introgressions. Between each introgression are *n*=100 simulated 10 kb sequences without introgression. Mean values over 20 consecutive regions for each *D*_*FOIL*_ statistic are shown by solid and dashed lines. Each of the eight inter-group introgressions (first eight non-shaded columns from left) all show a unique *D*_*FOIL*_ signature (i.e., a unique combination of +/-/0 signs for *D*_*FO*_, *D*_*IL*_, *D*_*FI*_, and *D*_*OL*_) consistent with Table 1. The ancestral introgressions (involving *P*_12_) can be distinguished between *P*_12_⇔*P*_3_ and *P*_12_⇔*P*_4_, but the direction cannot be determined by *D*_*FOIL*_ signature alone. For the intra-group introgressions (rightmost four non-shaded columns) and no-introgression treatments (narrow, shaded regions throughout), all three *D*_*FOIL*_ statistics are approximately 0 as expected.

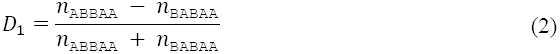

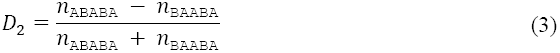

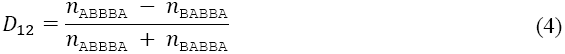

Positive values of *D*_1_ indicate *P*_2_⇔*P*_3_ introgression, while negative values indicate *P*_1_⇔*P*_3_ (Eaton and Ree 2013). Values not differing significantly from 0 indicate no introgression involving *P*_3_. For *D*_2_, positive and negative values indicate *P*_2_⇔*P*_4_ and *P*_1_⇔*P*_4_, respectively, while *D*_2_ = 0 indicates no introgressions involving *P*_4_. In addition, the *D*_12_ statistic was proposed as a means to determine the introgression donor and recipient taxa. *D*_12_ assumes that if both *P*_3_ and *P*_4_ exhibit the derived state, then the substitution must have occurred on the ancestral *P*_34_ branch (Fig. 1b). So *P*_3_ or *P*_4_ is assumed to be the introgression donor, and significant positive or negative values of *D*_12_ indicate the recipient is *P*_1_ or *P*_2_, respectively. Values not deviating significantly from 0 indicate *P*_1_ or *P*_2_ as the introgression donor and *P*_3_ or *P*_4_ as the recipient.

### Simulations

We tested the new *D*_*FOIL*_ statistics described below on a set of simulated chromosomes divided into 10 kb regions, each of which was allowed to evolve over a common phylogeny (Fig. 3b and 3c). All simulated chromosomes were generated using the Python libraries for the EggLib coalescent simulation engine (v2.1.7; De Mita and Siol 2012). A simulated population size of *N*_*e*_=10^6^ and per-base-per-generation mutation rate of *μ*=10^-9^ were used. Introgression was simulated by a temporary period of migration from the introgression donor population to the recipient population at a fixed rate (500 individuals per generation). Biallelic site patterns were counted for each region. To simulate the possible effects of convergent mutations in a finite sites mode, five additional substitutions were added randomly to each 10 kb region at sites with one or more existing derived states. The four *D*_*FOIL*_ and three Partitioned *D*-statistics were then calculated from the site pattern counts for each 10 kb window. Custom Python scripts were used for tabulation of site patterns and calculation of Partitioned *D*-statistics and *D*_*FOIL*_. Plots were generated with the *matplotlib* library (http://www.matplotlib.org). The simulation generation script and site count data sets are available through Dryad (URL PENDING). The DFOIL program used for all calculations and plots is for available for public use (http://www.bitbucket.org/jbpease/dfoil).

## Results

### The D_FOIL_ Test for a Symmetric Five-taxon Phylogeny

Our model describes a clade of five taxa connected by a symmetric phylogeny, denoted (((*P*_1_,*P*_2_),(*P*_3_,*P*_4_)),*O*), with the in-group taxa arranged in two sub-pairs (*P*_1_/*P*_2_ and *P*_3_/*P*_4_) and an out-group taxon (*O*; Fig. 1b). We define that *P*_3_ and *P*_4_ diverged (at time-before-present *T*_2_) before *P*_1_ and *P*_2_ (at *T*_3_), and also that the two sub-pair lineages diverged at *T*_1_. The labeling of the taxa (*P*_1_–*P*_4_) is arbitrary, as long as the sub-pairings are correct and the relationships between the three speciation times-before-present adhere to the relationships *T*_1_ > *T*_2_ ≥ *T*_3_ > 0.

As in the four-taxon case, we only sample mono- or biallelic sites (with the out-group always represented as A). 0 or 1 introgressions are allowed per region (at time-before-present *τ*). Reverse and convergent substitutions are expected to occur in negligible amounts, and are expected to affect all topologies equally (Durand et al. 2011). We refer to introgressions occurring between one taxon from each sub-pair as an inter-group introgression (of which there are *n* = 8 possible pairings; Fig. 3a) and those between taxa in the same sub-pair as an intragroup introgression (*n* = 4; Fig. 3a). Additionally, introgression between the ancestral branch *P*_12_ and *P*_3_ or *P*_4_ is possible when *T*_2_ > *τ* > *T*_3_, which will be referred to as an ancestral introgression (*n* = 4; Fig. 3a).

We propose a system of four *D*-statistics, collectively named *D*_*FOIL*_, to distinguish among the 16 possible introgressions in a symmetric five-taxon phylogeny. The name *D*_*FOIL*_ borrows from the “FOIL method,” a grade school mnemonic for multiplying two binomials (“First, Outer, Inner, Last”). We apply these labels to the four in-group taxa, and name the four *D*_*FOIL*_ statistics *D*_*FO*_ (“first”=*P*_1_/*P*_3_ *vs.* “outer” =*P*_1_/*P*_4_), *D*_*IL*_ (“inner”=*P*_2_/*P*_3_ *vs.* “last”=*P*_2_/*P*_4_), *D*_*FI*_ (“first” *vs.* “inner”), and *D*_*OL*_ (“outer” *vs.* “last”). These are defined as:

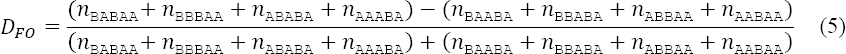

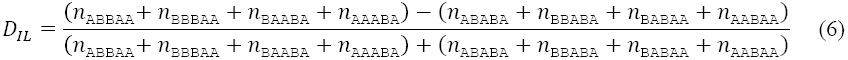

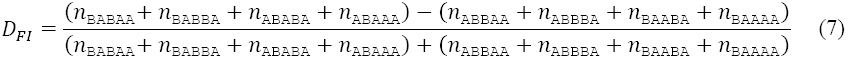

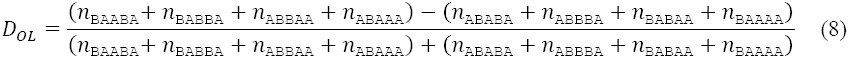

Each of these equations calculates the *D*-statistic for one of the four in-group taxa using the principles described previously. For example, *D*_*FO*_ tests *P*_1_ and describes the relative support for *P*_1_ being more closely related to *P*_3_ or *P*_4_ (i.e., the two taxa from the opposite sub-pair; Fig 1b). These two relationships are inferred by sampling two sets of gene trees (indicated by biallelic site patterns) that have an equal total probability of being sampled under the null hypothesis given the distribution of gene trees for a symmetric five-taxon phylogeny (“+*D*_*FO*_” and “–*D*_*FO*_” in Fig. 2b). Under ILS alone, the relative apparent strength of the two relationships represented on either side of the numerator of *D*_*FO*_ should be equal. Therefore, *P*_1_⇔*P*_3_ introgression will lead to more sampling of sites that support a closer relationship between *P*_1_ and *P*_3_, and therefore shift towards *D*_*FO*_ > 0. Alternatively, *P*_1_⇔*P*_4_ introgression will shift towards *D*_*FO*_ < 0. We also apply the principle of inverse patterns (e.g. BABBA and ABAAA), and so both terms of all *D*_*FOIL*_ tests include two pairs of inverse patterns. *D*_*IL*_, *D*_*FI*_, and *D*_*OL*_ are calculated identically to *D*_*FO*_, with *P*_2_, *P*_3_, and *P*_4_, respectively, as the focal taxon instead of *P*_1_, and *P*_1_/*P*_2_ used for comparison in *D*_*FI*_ and *D*_*OL*_. For all four tests, significant positive or negative values support a hypothesis of introgression for the focal taxon with one of the two taxa from the other sub-pair.

**FIGURE 1.**
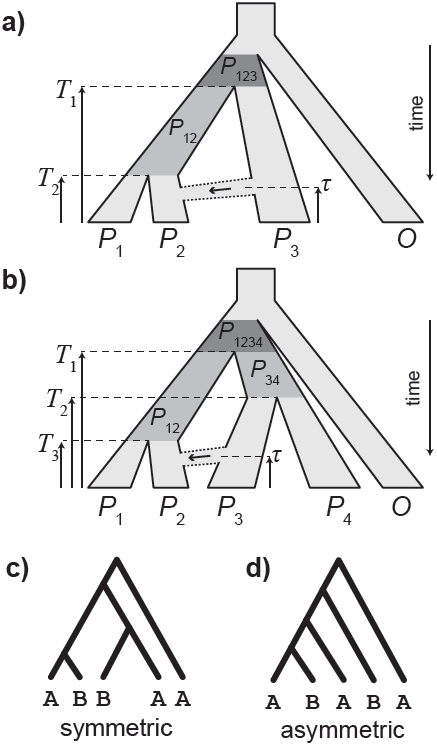
(a) A four-taxon phylogeny with three in-group taxa (*P*_1_-*P*_3_) and an out-group (*O*). (b) A five-taxon phylogeny with four in-group taxa (*P*_1_–*P*_4_) and an out-group (*O*). Ancestral branches are *P*_12_, *P*_34_, *P*_123_, and *P*_1234_. Introgressions (dotted region) are shown from *P*_3_⇒*P*_2_ in both (a) and (b). The time-before-present of the two or three speciations, respectively, are *T*_1_, *T*_2_, and *T*_3_, and time-since-introgression is *τ*. Five-taxon trees can have (c) symmetric or (d) asymmetric topologies, shown with examples of allelic state patterns ABBAA and ABABA, respectively.

Importantly, it should be noted that the constraint of equal probability for each of the two terms in the numerators of the *D*_FOIL_ statistics means that an analogous set of tests for an asymmetric five-taxon phylogeny cannot be constructed. In the asymmetric phylogeny ((((*P*_1_,*P*_2_),*P*_3_),*P*_4_)*O*) the relationship between *P*_1_ and *P*_3_ is not expected to be equal to *P*_1_ and *P*_4_ under the null. This occurs because a closer relationship between *P*_1_ and *P*_3_ only requires ILS to have occurred on one branch, while a closer relationship between *P*_1_ and *P*_4_ requires ILS on two ancestral branches. This highlights the special property that the four-taxon asymmetric and five-taxon symmetric phylogenies share with respect to ILS. In both topologies, all discordant coalescences must occur in the root, and therefore the exact probability distributions of discordant topologies are not needed to construct a test of the null hypothesis of no introgression.

### Direction of Introgression and Significance

As described thus far, the four *D*_*FOIL*_ statistics are simply individual applications of the *D*-statistic to each of the four in-group taxa. However, all four *D*_*FOIL*_ statistics considered collectively contain more information than the sum of the individual *D*-tests. In addition to identifying the taxa involved in introgression, the *D*_*FOIL*_ statistics can also provide information about the direction of inter-group introgressions, specifically identifying both the donor and recipient taxa. When an inter-group introgression occurs in a symmetric five-taxon phylogeny, the relationship changes not only between the donor and recipient taxa change, but also the recipient taxon and the donor’s sister taxon. For example, if *P*_3_⇒*P*_1_ occurs in an otherwise concordant gene tree, then the resultant topology becomes (((*P*_1_,*P*_3_),*P*_4_),*P*_2_),*O*) (Fig 2b, line viii). The shared history of *P*_3_ and *P*_4_ means that *P*_4_ also changes its relationship to *P*_1_. Conversely, if *P*_1_⇒*P*_3_ occurred the resultant topology is (((*P*_1_,*P*_3_),*P*_2_),*P*_4_),*O*) (Fig 2b, line vii), and now the relationship of *P*_2_ with *P*_3_ and *P*_4_ is altered by association. Even though both relationships change, the introgressing taxon should change more strongly than its sister taxon, and this disparity informs the overall assignment of donor and recipient. In this way, the signs (+, −, or 0) of the four *D*_*FOIL*_ tests collectively form a signature that provides information beyond the sum of its individual components. The *D*_*FOIL*_ tests individually indicate which taxa are introgressing, but collectively can also identify the donor and recipient taxa (or ancestral lineage) for a given introgression.

Each *D*_*FOIL*_ tests is separately assessed to be significantly positive, significantly negative, or not different than 0 by a two-tailed binomial test of the raw site counts for the left and right terms with a 50:50 expectation. Statistically significant positive and negative values are assigned a sign of “+” or “−,” while non-significant values are “0.” The four tests in the order (*D*_*FO*_, *D*_*IL*_, *D*_*FI*_, *D*_*OL*_) form the *D*_*FOIL*_ signature (Table 1). The appropriate *P*-value cutoffs for significance are determined by simulation, using the tested topology with no introgression. The 0.05 tail of simulated components of *D*_*FOIL*_ from regions with no introgression are used as the empirical *P*-value cutoffs. Individual *P*-values for each *D*_*FOIL*_ component can be calculated; however, it is expected that *D*_*FO*_ and *D*_*IL*_ will have approximately equal significance thresholds, as will *D*_*FI*_ and *D*_*OL*_. The difference in the thresholds between these two pairs of statistics should be approximately proportional to the difference in divergence times *T*_2_ and *T*_3_.

**TABLE 1.** The expected sign (+, -, or 0) of *D*_*FO*_, *D*_*IL*_, *D*_*FI*_, and *D*_*OL*_ for no introgression, eight inter-group introgressions, four ancestral introgressions, and two pairs of intra-group introgressions. Each inter-group introgression can be determined by a unique *D*_*FOIL*_ “signature.”

| **Introgression** | **$D_{FO}$** | **$D_{IL}$** | **$D_{FI}$** | **$D_{OL}$** |
| --- | --- | --- | --- | --- |
| None | 0 | 0 | 0 | 0 |
| $P_1 \Rightarrow P_3$ | + | + | + | 0 |
| $P_3 \Rightarrow P_1$ | + | 0 | + | + |
| $P_1 \Rightarrow P_4$ | − | − | 0 | + |
| $P_4 \Rightarrow P_1$ | − | 0 | + | + |
| $P_2 \Rightarrow P_3$ | + | + | − | 0 |
| $P_3 \Rightarrow P_2$ | 0 | + | − | − |
| $P_2 \Rightarrow P_4$ | − | − | 0 | − |
| $P_4 \Rightarrow P_2$ | 0 | − | − | − |
| $P_{12} \Rightarrow P_3$ | + | + | 0 | 0 |
| $P_3 \Rightarrow P_{12}$ | + | + | 0 | 0 |
| $P_{12} \Rightarrow P_4$ | − | − | 0 | 0 |
| $P_4 \Rightarrow P_{12}$ | − | − | 0 | 0 |
| $P_1 \Leftrightarrow P_2$ | 0 | 0 | 0 | 0 |
| $P_3 \Leftrightarrow P_4$ | 0 | 0 | 0 | 0 |

### Application and accuracy of the D_FOIL_ method

To demonstrate the effectiveness of the *D*_*FOIL*_ method, we simulated the evolution of chromosomes over a common phylogeny (Fig. 3a and 3b). Along the chromosome, we simulated sixteen sets of *n*=500 loci of length 10 kb for each of the sixteen possible introgressions (eight inter-group, four ancestral branch, four intra-group). Between each introgressed region, *n*=100 regions of no-introgression control were included for visual contrast (Fig. 3d, shaded). In all sixteen cases and the control, the *D*_*FOIL*_ signature (signs of the four *D*_*FOIL*_ tests) that was observed in the simulations matched the theoretical expectations (Fig. 3d and Table 1). For example, the introgression *P*_1_⇒*P*_3_ (Fig. 3d, leftmost region) shows positive values for *D*_*FO*_, *D*_*IL*_, and *D*_*FI*_, while *D*_*OL*_ ≈ 0, in agreement with the expected +/+/+/0 signature. In each of the eight inter-group introgressions (Fig. 3d, leftmost eight regions), a unique *D*_*FOIL*_ signature clearly distinguishes each case. Ancestral introgressions involving branch *P*_12_ (Fig. 3c and middle of 3d) were distinguishable from cases of introgression with *P*_3_ or *P*_4_, but the direction of these introgression cannot be determined by the *D*_*FOIL*_ signature alone (see Discussion). As expected, the average value for all four *D*_*FOIL*_ tests in intra-group introgressions (Fig. 3d, rightmost four regions) is 0. Therefore, the *D*_*FOIL*_ tests can distinguish the taxa involved in inter-group and ancestral introgressions, and additionally the direction of introgression for inter-group introgressions.

### The Five-Taxon “Partitioned” D-statistics

We also used the *D*-statistic framework described above to examine the Partitioned *D*-statistics (Eq. 2-4; Eaton and Ree 2013). As previously noted, inverse pairs of biallelic patterns (e.g. ABBAA/BAABA) both indicate the same underlying gene tree when introgression is considered. This means that the site pattern counts used in *D*_1_ and *D*_2_ of the Partitioned *D*-statistic are not necessarily unique to the introgressions they propose to test. Specifically, the left term of *D*_1_ (*n*_ABBAA_) and right term of *D*_2_ (*n*_BAABA_) are directly related, as are the right term of *D*_1_ (*n*_BABAA_) and left term of *D*_2_ (*n*_ABABA_). Therefore, an inter-group introgression between any two taxa should change both *D*_1_ and *D*_2_ in opposite directions.

This inverse relationship between *D*_1_ and *D*_2_ means that introgression between one pair of taxa creates a “mirror effect” that makes the other pair of taxa falsely exhibit evidence of introgression. Therefore, while *D*_1_ and *D*_2_ in this form can detect the occurrence of an intergroup introgression, the mirror effect makes it unclear which two taxa are actually exchanging alleles. This effect is a consequence of using biallelic patterns where two (out of four) taxa exclusively share one state, since, by default, the other two taxa share the opposite state.

In all cases of inter-group introgression, the simulation data showed an inverse relationship between *D*_1_ and *D*_2_, confirming the mirror effect (Fig. 4). For example, the introgressions *P*_1_⇒*P*_3_ and *P*_3_⇒*P*_1_ (Fig. 4, left side) are expected to be *D*_1_ < 0 and *D*_2_ ≈ 0. However, in both cases *D*_2_ showed positive values that mirrored the negative values of *D*_1_, making it falsely seem as though *P*_2_⇔*P*_4_ was also occurring in both cases. A similar result was observed for the introgressions *P*_1_⇒*P*_4_ and *P*_4_⇒*P*_1_ (Fig. 4, right side). In both cases, the expected value of *D*_1_ is 0, but instead *D*_1_ mirrors *D*_2_, with positive values of *D*_1_ that make it appear as though *P*_2_⇔*P*_3_ has also occurred. We can conclude both from theory and simulations that *D*_1_ and *D*_2_ are inversely related, and therefore are non-specific in determining which pair of taxa have introgressed.

The patterns counted by *D*_12_ could also be arrived at through introgressions other than the ones intended to be tested. For example, ABBBA is assumed by Eaton and Ree (2013) to be the result of a *P*_3_⇒*P*_2_ or *P*_4_⇒*P*_2_ transfer of the B state from a pre-introgression pattern of AABBA. However, ABBBA can also be formed from the patterns ABABA and ABBAA by *P*_2_⇒*P*_3_ or *P*_2_⇒*P*_4_, respectively. This means that for any given inter-group pair of taxa, introgressions in both directions will both raise or both lower the value of *D*_12_. This occurs because Eaton and Ree (2013) assume that the substitution A→B occurred on the ancestral *P*_34_ branch, which also forces *P*_3_ and *P*_4_ to both be state B. This effectively collapses *P*_3_ and *P*_4_ to a single branch and makes *D*_12_ a four-taxon *D*-statistic.

Simulations also confirmed these conclusions, showing that *D*_12_ values for a given pair of taxa are either positive or negative regardless of direction of introgression. For example, among the four introgressions shown in Figure 4, *P*_3_⇒*P*_1_ and *P*_4_⇒*P*_1_ exhibit negative values of *D*_12_, respectively, as expected. However, *P*_2_⇒*P*_3_ and *P*_2_⇒*P*_4_ are expected to be *D*_12_ ≈ 0, but while they appear closer to zero, both cases still show a non-zero value of *D*_12_ (approximately at the same level as *D*_1_) with the same sign as introgression in the opposite direction. These simulations confirm that *D*_12_ will have the same sign for an introgression between given pairs of taxa regardless of direction.

**FIGURE 4.**
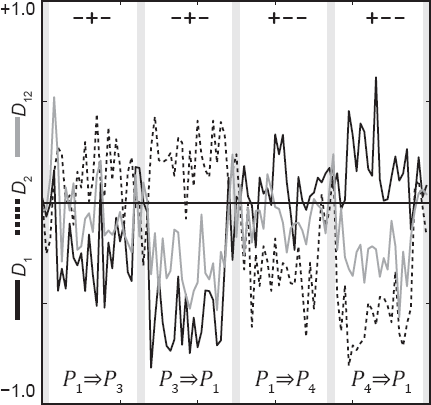
Partitioned *D*-statistics calculated for the same simulated chromosome and phylogeny as shown in Figure 3b. The Partitioned *D*-statistics suffer from a “mirror effect” due to the inverse relationship of *D*_1_ and *D*_2_. For example, in the region under introgression *P*_1_⇒*P*_3_ (leftmost column), *D*_1_ is negative as expected, but *D*_2_ is positive (and should =0). Also, *D*_12_ has the same sign for introgressions in both directions for each pair of taxa.

## Discussion

### Robustness of the D_FOIL_ Tests to Special Cases

There are several special cases in which introgression occurs at particular points in the phylogeny and for which detection of these events might be more problematic. However, the triangulation approach of the *D*_*FOIL*_ tests allows them to still function even in these special cases. If speciation of the two subgroups occurred at approximately the same time (i.e., *T*_2_ ≈ *T*_3_), such that the difference between the two is not clearly discernible, the signs of all four statistics would remain unaffected. This would, however, negate the possibility of ancestral introgression, since both speciation events occur at approximately the same time.

If introgression occurs very close to the second speciation event (i.e., *τ* ≈ *T*_3_) this would cause *D*_*FI*_ and *D*_*OL*_ to reduce to 0 in the cases where introgression is *P*_1_/*P*_2_⇒*P*_3_/*P*_4_. In this instance, *D*_*FI*_ and *D*_*OL*_ are only reflecting the reality that the introgression involves a population of *P*_1_ or *P*_2_ that is so recently diverged that they are practically indistinguishable genetically. So as *τ* approaches *T*_3_, the *D*_*FOIL*_ statistics effectively convert to the signature of an ancestral introgression (+/+/0/0 or -/-/0/0).

One assumption that may be violated in real datasets is the expectation of uniform substitution rates across the tree. If a particular taxon has a much overall higher substitution rate than the others, then all distances involving that taxon would be relatively higher due to an increased rate of substitution rather than an introgression involving its sister taxon. There are two straightforward solutions to these lineage-specific effects. The first would be to weight the expected *D*-statistic values away from 50:50 when there is a prior expectation that substitution rates are not equal between sister taxa in a subgroup. The second would be to simply exclude the terminal-branch-substitution site patterns (AAABA, AABAA, ABAAA, and BAAAA) from all calculations. As long as these counts are excluded from both sides of the equation, the expectation of equality of the left and right terms of each *D*_*FOIL*_ statistic is not violated.

### Ancestral Introgressions

Ancestral introgressions can also be detected by the *D*_*FOIL*_ framework, though the direction of this introgression cannot be detected strictly by the *D*_*FOIL*_ signature. When calculating a single, genome-wide estimate for each of the *D*_*FOIL*_ tests, determining the direction of ancestral introgressions is not possible. However, if, for example, both *P*_12_⇒*P*_3_ and *P*_3_⇒*P*_12_ are present at different locations in the genome, the difference in their expected values for *D*_*FO*_ and *D*_*IL*_ could be used to distinguish between these two cases in the different regions.

### More Taxa, More Models

In order to detect introgression in a phylogeny of six or more taxa, the simplest option is simply to subsample a part of the tree in the appropriate configuration and either to use the four-taxon *D*-statistic or the five-taxon *D*_*FOIL*_ tests. The *D*_*FOIL*_ tests offer the added feature of information about the direction of introgression in the subgroup of interest, when a five-taxon subset is available in the symmetric configuration. Aside from the subsampling option, we envision that a formal expansion of the model for more than five taxa would also be possible for certain tree topologies using the previously described principles of the *D*-statistic. The symmetric five-taxon tree represents a special case with respect to ILS, since the topology dictates that all discordant coalescences must occur in the root. This leads to a simple distribution of gene trees and provides a topology where each of the four in-group taxa can be compared in a straightforward manner with the two taxa in the opposite sub-pairs. Beyond this special case though, the probability distribution of gene trees becomes far more complex due to increasing topological constraints (Rosenberg 2002; Degnan and Salter 2005; Degnan and Rosenberg 2006). Therefore, while an explicit solution in larger phylogenies is not impossible, the more straightforward solution is simply to subsample four taxa of interest and an appropriate outgroup.

Some care must be taken when subsampling from a larger tree, or when planning which taxa to sample from nature. Introgression from taxa not included in the sample—what Durand et al. (2011) refer to as “ghost taxa”—could cause incorrect results using *D*_*FOIL*_ or the *D*-statistic. When a pair of taxa introgress within the sampled clade, we infer introgression from closer relationships between these two taxa. However, if we only sampled the recipient of introgressed alleles, this taxon will appear to be unusually divergent from its sister taxon. Alternatively, a false positive of introgression could be detected if the ghost taxon has shared ancestry with one of the sampled taxa, or two taxa in the sample were recipients of introgressed alleles from the same ghost taxon. In general, however, we would expect that introgressions from taxa not in the sample would simply result in a noisier signal of introgression or increased rate of false negatives.

In the *D*_*FOIL*_ tests, only biallelic site patterns are used to determine the phylogenetic relationships between taxa. This simple distance measure is particularly ideal for closely related species with few sequence differences and many biallelic sites. Since the site patterns used in *D*_*FOIL*_ fundamentally derive from relative phylogenetic relationships and gene trees, these same underlying phylogenetic relationships could be used to build a *D*_*FOIL*_ method using other models of sequence evolution. The conceptual framework would remain the same, only the model of inferring phylogenetic relationships between taxa would be altered.

### Conclusions

In clades of closely related species, where ILS is prevalent or there are few sequence differences (or both), it can be difficult to detect introgression. The *D*_*FOIL*_ system offers a simple means to infer introgression in a symmetric five-taxon clade, requires little computational power, and functions even with relatively little sequence divergence and high levels of ILS. When computed on multiple alignments of whole-chromosomes, the added spatial context will allow for the detection of localized introgressions throughout the genome. Spatial context also makes possible the detection of introgressions among different combinations of taxa at various locations throughout the genome. The *D*_*FOIL*_ statistic is also designed specifically to detect the direction of introgression, when sequence data for sufficient taxa are available.

The *D*_*FOIL*_ tests can be used to test for general introgression across the genome in datasets without reference genomes, using RNA-Seq, RAD-Seq, or other targeted sequencing technologies. Since such loci may not have a known order along chromosomes, these data are not suitable for locus-by-locus testing of introgression. For these data, a single genomic mean value for each of the four *D*_*FOIL*_ tests can be computed (as was done by Green et al. 2010), and the average direction of introgression can also be determined. This implementation of *D*_*FOIL*_ offers a more diffuse — but still informative — look at introgression from a smaller subset of data without spatial context.

## Supplementary Material

Simulation scripts, simulated site pattern counts and D-statistics are available through Dryad (URL PENDING). The DFOIL script is available on BitBucket (http://www.bitbucket.org/dfoil)

## Acknowledgements

We thank Julien Dutheil, Michael Fontaine, Daniel Neafsey, and Nora Besansky for helpful discussion, and Gregg Thomas for comments on the manuscript.

